# Dual Inhibition of CDK4/6 and mTORC1 Establishes a Preclinical Strategy for Translocation Renal Cell Carcinoma

**DOI:** 10.1101/2025.07.11.663903

**Authors:** Shikha Gupta, Prateek Khanna, Eddy Saad, Renee Maria Saliby, Shatha AbuHammad, Jiao Li, Bingchen Li, Prathyusha Konda, Usman Ali Ahmed, Ananthan Sadagopan, Qingru Xu, Ziad Bakouny, Wenxin Xu, Ramaprasad Srinivasan, Toni K. Choueiri, Srinivas R. Viswanathan

## Abstract

**Purpose:** Translocation renal cell carcinoma (tRCC) is a rare and aggressive subtype of kidney cancer driven by an oncogenic fusion involving a transcription factor in the *MiT/TFE* gene family, most commonly *TFE3*. Treatment of tRCC currently lacks a clear standard of care, underscoring the pressing need to nominate new therapeutic targets with mechanistic rationale in this cancer.

**Experimental Design:** In this study, we applied integrative genomic approaches to identify activation of the cyclin-dependent kinase 4/6 (CDK4/6) and mammalian target of rapamycin complex 1 (mTORC1) pathways in tRCC. We tested the activity of CDK4/6 inhibitors (CDK4/6i), alone or in combination with mTORC1-selective inhibition, using *in vitro* and *in vivo* models of tRCC.

**Results:** tRCC tumors displayed multiple genomic and transcriptional features associated with activation of the CDK4/6 and mTORC1 signaling pathways. Genetic or pharmacologic inhibition of CDK4/6 suppressed tRCC cell growth and induced cell cycle arrest *in vitro* but was not cytotoxic, with rapid cell regrowth observed after drug withdrawal. The mTORC1-selective inhibitor, RMC-5552, potently reduced translation of Cyclin D1, which complexes with CDK4/6 proteins to regulate G1-S cell cycle progression. Combined treatment with the CDK4/6 inhibitor, palbociclib, and RMC-5552 resulted in synergistic suppression of tRCC cell viability and increased markers of apoptosis *in vitro*. The combination of palbociclib and RMC-5552 in a tRCC xenograft model showed greater efficacy than either single agent while also being well-tolerated.

**Conclusions:** Our study indicates the therapeutic potential of combined CDK4/6 and mTORC1 inhibition in tRCC, providing the rationale for further clinical evaluation of this strategy.

**Translational Relevance:** Translocation renal cell carcinoma (tRCC) is a rare and aggressive form of kidney cancer that accounts for 2-5% of all RCCs in adults and around 50% of RCCs in children. tRCC lacks effective therapies and represents a significant unmet clinical need. Therapies approved for clear cell RCC (ccRCC), including VEGF/multikinase inhibitors and immune checkpoint blockade, typically show decreased efficacy in tRCC. Via an integrative genomic analysis, we nominate combined inhibition of the cyclin-dependent kinase 4/6 (CDK4/6) and mammalian target of rapamycin complex 1 (mTORC1) pathways in tRCC. We demonstrate that combining the CDK4/6 inhibitor palbociclib with the mTORC1-selective inhibitor RMC-5552 synergistically inhibits the growth of tRCC cells *in vitro* and of tRCC xenografts *in vivo*, where the combination is well-tolerated. This proof-of-concept study provides preclinical evidence supporting CDK4/6 inhibitor-based combination regimens tailored to tRCC biology, offering a foundation for future clinical studies in tRCC, which currently has no clear standard of care.

## Introduction

Translocation renal cell carcinoma (tRCC) is a highly aggressive neoplasm that is defined by a gene fusion involving a transcription factor in the *MiT/TFE* gene family, most commonly *TFE3* (1). tRCC comprises approximately 2-5% of all RCCs in adults and around 50% of RCCs in children (1,2); however, due to its histologic similarities with other subtypes of kidney cancer and frequent misclassification, the true incidence of tRCC is likely higher (3).

tRCC is clinically aggressive with no effective therapies for metastatic disease and a median survival of less than 2 years (4–6). Approved agents for clear-cell RCC (ccRCC, the most common RCC subtype), such as VEGF/multikinase inhibitors and immune checkpoint blockade, have not been specifically tested in tRCC. Although these treatments are often employed empirically in tRCC, they typically yield poorer response rates than in ccRCC (4,5,7–9). Therefore, new molecular targets in tRCC, which are based on strong mechanistic rationale and linked to its unique biology, are urgently needed.

Recent studies have begun to elucidate the genomic and transcriptomic features of tRCC (10–15). Although there are few recurrent alterations in tRCC apart from the driver fusion, inactivation of the *CDKN2A* gene (which encodes p16^INK4A^, an endogenous negative regulator of CDK4/6), occurs in about 20% of cases (12,15). Other studies have identified a proliferative gene signature in tRCC, typified by high expression of cell cycle regulators including *CDK4, CDK6, and E2F* target genes (14,16). Additionally, increased expression of proteins involved in cell cycle progression has also been noted in tRCC (13). Altogether, these data nominate cell cycle dysregulation as a potential therapeutic target in tRCC.

CDK4 and CDK6 form complexes with D-type cyclins (Cyclin D1, D2, and D3) to regulate the G1 to S phase transition of the cell cycle. The CyclinD-CDK4/6 complex phosphorylates the retinoblastoma (Rb) protein, leading to the release of E2F transcription factors that promote S-phase entry (17). Therapeutically, this pathway can be targeted in cancer with CDK4/6 inhibitors (CDK4/6i), of which three have been FDA approved for the treatment of hormone receptor-positive metastatic breast cancer: palbociclib, abemaciclib, and ribociclib (18,19). These drugs have transformed the treatment of hormone receptor-positive breast cancer, where they are used in combination with aromatase inhibitors or estrogen receptor (ER) antagonists. Notably, inhibition of the ER axis works in concert with CDK4/6i by suppressing levels of *CCND1* (a direct gene target of ER) (18). However, resistance to CDK4/6i has been a major barrier to the broader deployment of these agents, and multiple mechanisms have been described including alterations in the cell cycle pathway (including Rb and Cyclin D alterations), activation of growth factor signaling, and metabolic reprogramming, among others (20).

In this study, we explored the potential of CDK4/6 inhibition as a therapeutic approach in tRCC and tested CDK4/6i in combination with RMC-5552, a recently characterized mTORC1-selective inhibitor that is currently in clinical trials (21–24). Given the ability of RMC-5552 to potently suppress phosphorylation of 4-EBP1 and thereby lead to reduced levels of Cyclin D1 protein (24), we reasoned that the combination of CDK4/6i and RMC-5552 may display therapeutic efficacy in tRCC.

## Results

### CDK4/6 and mTORC1 signaling are activated in tRCC and are regulated by TFE3 fusions

Prior studies have nominated several molecular pathways that may be activated in tRCC (e.g. PI3K/mTOR, MET, RET, WNT), although these findings have not yet been translated clinically (25–30). Additionally, several lines of evidence suggest that the CDK4/6 pathway may be activated in tRCC, through both genomic mechanisms (e.g. loss of *CDKN2A*) (12,15) and transcriptional mechanisms (e.g. upregulation of genes involved in cell cycle progression) (14,16).

We hypothesized that activation of the CDK4/6 pathway in tRCC could be analogous to the setting of hormone receptor positive breast cancer, with both CDK4/6 expression and cyclin D expression directly linked to the driver TFE3 fusion (**Fig. 1A**) (18). We compared transcriptional profiles between tRCC tumors and adjacent normal tissues in two recently published tRCC cohorts profiled by RNA-Seq (31,32). In both datasets, we observed a significant enrichment of both E2F targets (indicative of activated CDK4/6 signaling) and mTORC1 signaling in tRCC tumors relative to normal tissue (**Fig. 1B-C and S1A-B**). Consistent with these findings, analysis of expression of CDK4/6 and Cyclin D genes revealed higher expression in tRCC vs. normal samples (**Fig. 1D**). This was confirmed in several cell line models of tRCC (**Fig. S1C**). Interestingly, E2F activation and high expression of CDK4/6 as well as D-type Cyclins appeared to be independent of *CDKN2A* alteration status (**Fig. S1C**). Notably, recurrent alterations in Rb have not been reported in tRCC, suggesting that tRCCs would not harbor this intrinsic resistance mechanism to CDK4/6 pathway inhibition (12,31–34). We also observed that high expression of E2F target genes was associated with inferior survival in tRCC (n=25 tRCC, n=16 normal), even after controlling for differences in stage (log-rank p=0.0022, **Fig. 1E**); this may suggest inter-patient differences in the level of E2F signaling driven by the fusion. Taken together, these analyses point to activation of the CDK4/6 pathway in a majority of tRCC cases and suggest that targeting cell cycle progression might be of therapeutic benefit in this cancer.

**Fig 1:**
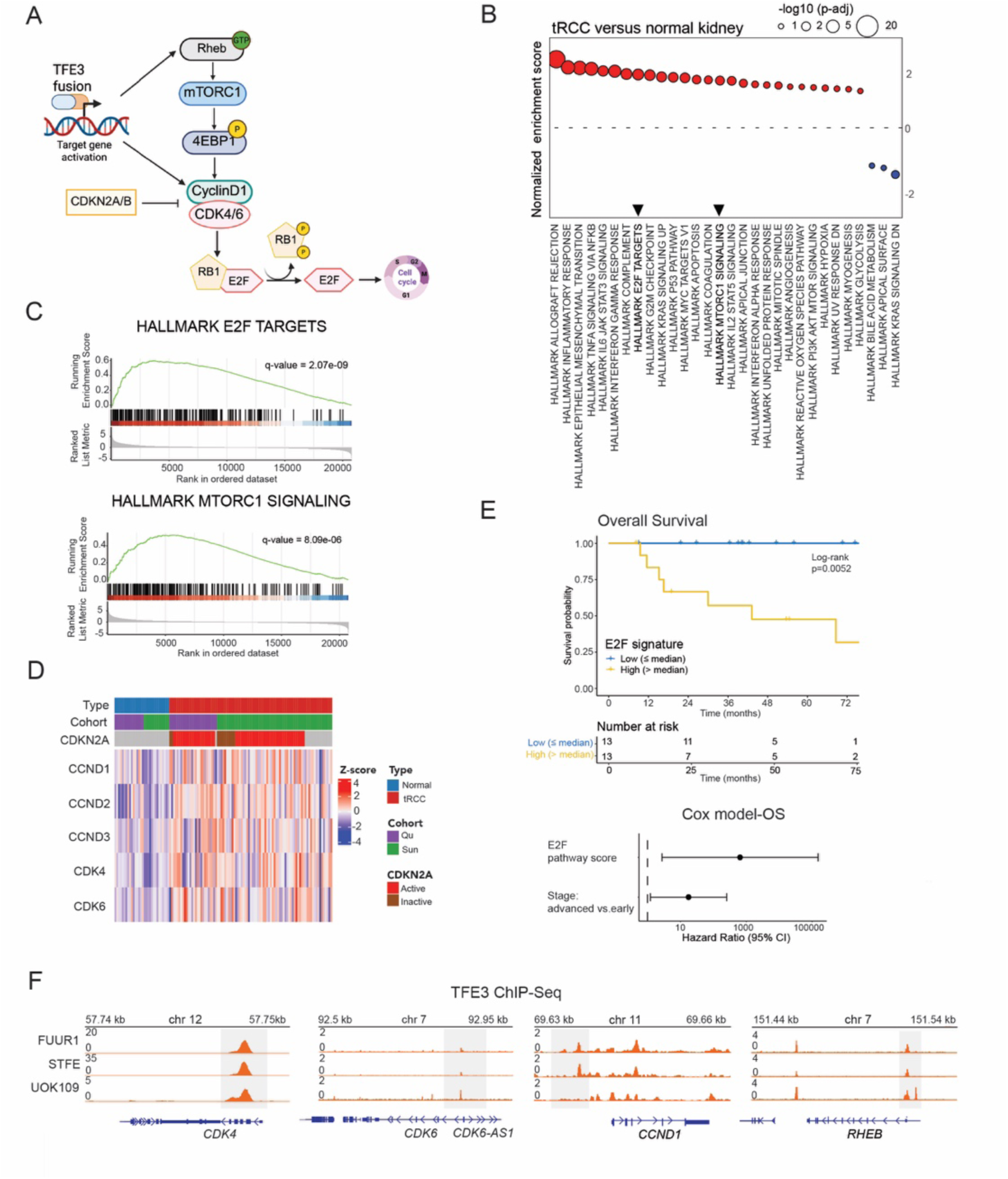
Activation of CDK4/6 and mTORC1 pathways in tRCC. (A) Schematic depicting activation CDK4/6 and mTORC1 signaling linked to the driver TFE3 fusion in tRCC. (B) Gene set enrichment analysis (GSEA) for tRCC versus adjacent normal kidney samples in *Sun et al*., 2021 (14), highlighting the top 30 enriched pathways (n= 63 tRCC, n= 14 normal) (C) GSEA enrichment plot for E2F targets (top) and mTORC1 signaling (bottom) from analysis in (B) from *Sun et al*., 2021 (14). (D) Heatmap depicting gene expression for *CDK4, CDK6,* and D-type Cyclins in tRCC versus normal samples. Data obtained from *Sun et al., 2021, Qu et al., 2022,* (n= 88 tRCC, n= 30 normal) (13,14). (E) Kaplan Meier survival plot for tRCC patients from *Qu et al., 2022* (13), stratified based on median-dichotomized *E2F* signature score, p value by log-rank test (top); forest plot for hazard ratio for overall survival with multivariable Cox model using E2F pathway score and early or advanced disease stage (bottom). (F) IGV tracks for TFE3 fusion ChIP-seq peaks from *Li et al., 2025* (35) at the *CDK4*, *CDK6*, *CCND1* and *RHEB* loci across three tRCC cell lines.

To determine whether activation of E2F and mTORC1 pathways was linked to the TFE3 fusion, an oncogenic transcription factor, we interrogated our recently published TFE3 ChIP-Seq data for selected genes involved in CDK4/6 and mTORC1 signaling (35). We observed strong TFE3 fusion peaks upstream of *CDK4*, *CDK6* and *CCND1* in multiple tRCC cell lines. Additionally, we observed a *TFE3* peak upstream of *RHEB*, an activator of mTORC1 in its GTP-bound form (36) (**Fig.1F**). Collectively, these data point to both CDK4/6 activation and mTORC1 signaling upregulation as defining features of tRCC, transcriptionally driven by the TFE3 fusion.

### tRCC cells are reversibly sensitive to CDK4/6 inhibition

Given transcriptional and proteomic evidence supporting the activation of CDK4/6 signaling in tRCC, we next assessed whether *CDK4* and *CDK6* represent genetic dependencies in this cancer type. We interrogated genome-scale CRISPR (clustered regularly interspaced short palindromic repeats) screening data from three tRCC cell lines (35,37) and found that *CDK4* was a strong dependency in all three tRCC cell lines, while *CDK6* was a strong dependency only in UOK109 cells (**Fig. 2A**).

**Fig 2:**
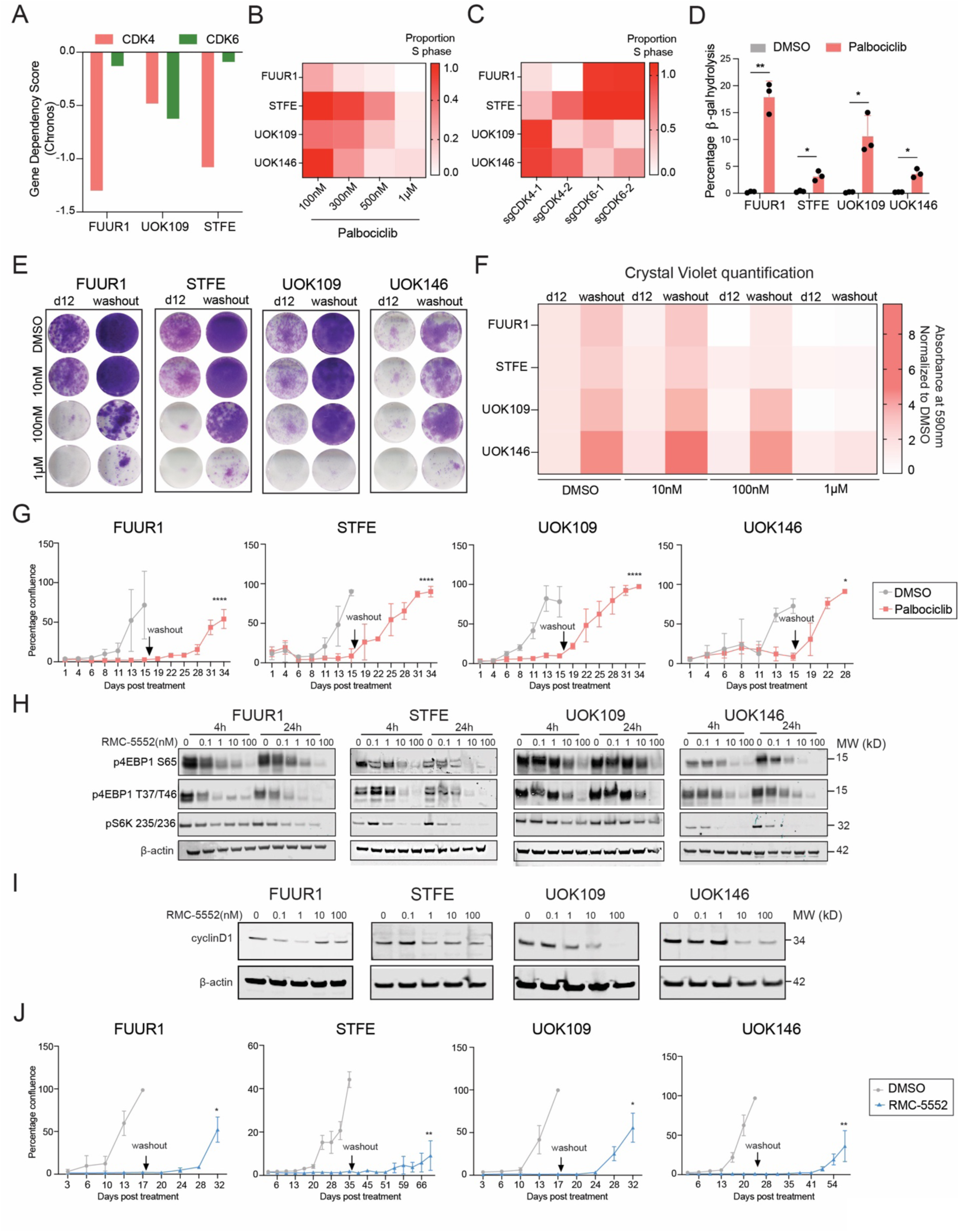
tRCC cells are reversibly sensitive to CDK4/6 inhibition and show reduced cyclinD1 expression upon suppression of mTORC1 signaling. (A) Gene dependency score (Chronos) for *CDK4* and *CDK6* in FUUR1, STFE and UOK109 cells in genome-scale CRISPR screening performed in a separate study (37). (B) Heatmap depicting the proportion of cells in S phase normalized to control (DMSO) at different doses of palbociclib (range 100nM-1mM) after 72 hours of treatment across 4 tRCC cell lines. Data representative of n=3 biological replicates per cell line. (C) Heatmap depicting the proportion of cells normalized to non-targeting control, in S phase upon genetic knockout of *CDK4* and *CDK6* using two different guide RNAs for each, 7 days post selection. Data representative of n=3 biological replicates per cell line. (D) Senescence-associated β-galactosidase activity after treatment with 1µM palbociclib for 72 hours, as determined by flow cytometry. Data representative of n=3 biological replicates per cell line, unpaired t-test *p<0.05*, p<0.01\*\**. (E) Colony forming assay for tRCC cells treated with 10nM, 100nM and 1µM of palbociclib after 12 days of drug treatment with DMSO vehicle (left column) or after 12 days of treatment followed by 12 days of washout (right column). Images for one of the three representative biological replicates shown. (F) Quantification of colony formation assay from mean of three independent biological replicates after dissolution of crystal violet staining. Data plotted as normalized absorbance relative to DMSO, n=3 biological replicates per cell line, unpaired t-test *p<0.05*, p<0.01\*\**. (G) Cell confluence over time in tRCC cells treated with DMSO (control) or palbociclib (300nM) with drug replenishment every 3rd day. Palbociclib treated cells were subjected to drug washout when the control cells crossed 75% confluency and confluence continued to be monitored in both arms, data representative of n=3 biological replicates per cell line, 2-way *ANOVA p<0.05*, p<0.001\*\*\*\** on last day of the experiment. (H) Immunoblot analysis for phospho-4EBP1 S65, phospho-4EBP1 T37/T46, phosphor-S6K 235/236, 4 hours and 24 hours post RMC-5552 treatment from 0.1-100nM dose range across tRCC cell lines, data representative of n=3 biological replicates per cell line. (I) Immunoblot analysis for CyclinD1 after 72 hours of RMC-5552 treatment from 0.1nM-100nM dose range across tRCC cell lines, data representative of n=3 biological replicates per cell line. (J) Cell confluence over time in tRCC cells treated with DMSO (control) or RMC-5552 (200nM) with drug replenishment every 3rd day. RMC-5552 treated cells were subjected to drug washout when the control cells crossed 75% confluency and confluence continued to be monitored in both arms, data representative of n=3 biological replicates per cell line, 2-way *ANOVA p<0.05*, p<0.01\*\**.

The clinically-approved CDK4/6 inhibitors have different levels of potency against CDK4 and CDK6 as well as varying spectra of activity against other kinases. Given our finding that both CDK4 and CDK6 may represent genetic dependencies in tRCC, we selected palbociclib as the primary CDK4/6i for further testing, as it has comparable affinity for the CDK4/Cyclin D and CDK6/Cyclin D complexes (whereas abemaciclib and ribociclib are more CDK4-selective) (38). We verified that palbociclib treatment suppresses growth across a panel of tRCC cell lines harboring different TFE3 fusions (FUUR1: *ASPSCR1-TFE3*; *sTFE*: *ASPSCR1-TFE3*; UOK109: *NONO-TFE3*; UOK146: *PRCC-TFE3*) (**Fig. S2A**). In addition, phosphorylation levels of RB1 decreased in all cell lines in a dose-dependent manner, validating on-target activity of palbociclib on the CDK4/6 complex (**Fig. S2B**). Moreover, treatment with palbociclib was associated with an increase in cell size (**Fig. S2C**) and an accumulation of cells in G1 with reduced S phase, consistent with drug-induced G1/S arrest (**Fig. 2B and Fig. S2D**).

To validate these findings genetically, we generated individual CRISPR knockouts for *CDK4* or *CDK6* using two different sgRNAs for each target (**Fig. S2E**). Consistent with the pharmacologic observations from palbociclib mediated CDK4/6 inhibition, the proportion of tRCC cells with either *CDK4* or *CDK6* knockout showed a significant reduction in the S phase population compared to the control (**Fig. 2C**). Interestingly, however, FUUR1 and sTFE cells appeared more dependent on *CDK4* knockout while UOK109 and UOK146 cells appeared more dependent on *CDK6* knockout, supporting our rationale to prioritize palbociclib as a CDK4/6i with potent activity against both CDK4 and CDK6.

In other cancer types, CDK4/6i have been shown to induce senescence following G1/S arrest (39). In line with this, we observed that treatment of tRCC cell lines with palbociclib resulted in an increase of the senescence-associated marker β-galactosidase (**Fig. 2D**). However, this phenotype was reversible, and washout of palbociclib in both a colony formation assay (**Fig. 2E, F**) and long-term culture assay (**Fig. 2G**) led to rapid cell regrowth. This suggests that tRCC cells are sensitive to palbociclib-induced growth arrest, but that single-agent treatment is not cytotoxic.

### mTORC1 inhibition suppresses cyclinD1 protein expression in tRCC

Multiple prior studies have implicated mTORC1 signaling as a resistance mechanism to CDK4/6 inhibition, providing the rationale for combining mTOR inhibition with CDK4/6i clinically (40–44). However, pan-mTOR (mTORC1/mTORC2) inhibitors are limited by toxicity, hyperglycemia, and the release of AKT-dependent feedback inhibition on receptor tyrosine kinases – effects chiefly attributable to mTORC2 suppression (45,46). In addition, pan-mTOR inhibitors only modestly suppress the phosphorylation of 4EBP1, which is critical for the regulation of Cyclin D expression (24). Recently, a class of bi-steric compounds – typified by RMC-5552 – has been described: these compounds have high selectivity for mTORC1 over mTORC2, potently suppress 4EBP1 phosphorylation and do not cause hyperglycemia (24).

We first tested the single-agent activity of RMC-5552 in tRCC cells and observed that it potently suppressed viability across our cell line panel (**Fig. S2F**). Immunoblot analysis revealed that inhibition of mTORC1 using RMC-5552 does not markedly suppress phospho-Akt levels (**Fig. S2G**), in contrast with pan-mTOR inhibitors that typically suppress p-AKT (47). We then tested the activation of 4EBP1 and S6K, downstream effectors of mTORC1, by measuring their phosphorylation levels. Indeed, we observed a pronounced reduction in p4EBP1 and pS6K levels as soon as 4 hours of RMC-5552 treatment within 1-10nM range (**Fig. 2H**) highlighting the potency of this compound in selectively inhibiting mTORC1 activity. This reduction in p-4EBP1 was accompanied by a corresponding decrease of cyclin D protein levels in all cell lines (**Fig. 2I**). However, similar to our results with palbociclib monotherapy, a long-term culture assay revealed that, while RMC-5552 suppresses cell proliferation with continuous drug exposure, cells regrow soon after drug washout (**Fig. 2J**). Together, these results indicate activity to both CDK4/6i and mTORC1 inhibition in tRCC but cell regrowth after washout is reversible in both settings.

### Combined inhibition of CDK4/6 and mTORC1 signaling inhibits tumor growth *in vivo*

We reasoned that the combination of CDK4/6i and mTORC1 inhibition could be efficacious in tRCC owing to coordinate suppression of both CDK4/6 kinase activity and Cyclin D protein levels, resulting in decreased activity of the CDK4/6-Cyclin D complex. Such an approach has been clinically effective in hormone-receptor positive breast cancer, where CDK4/6i are combined with ER pathway inhibitors, which decrease Cyclin D levels (48–50).

We pre-treated tRCC cell *lines in* vitro with palbociclib for 48 hours followed by a combination of palbociclib and RMC-5552 (0-10μM both) for 72 hours and quantified cell number using DAPI staining of nuclei. We observed that combination treatment with palbociclib and RMC-5552 suppressed viability more than either agent alone, with evidence of synergy observed to varying degrees, and at similar dose combinations, across the four tRCC cell lines tested (**Fig. 3A, Fig. S3A**). Interestingly, an increase in forward scatter profile – suggestive of G1 arrest – was observed with palbociclib monotreatment but not with RMC-5552 or combination treatment (**Fig. S3B**). Consistent with this finding, we observed no significant induction of β-galactosidase by RMC-5552 alone and a reduced level with combination treatment relative to palbociclib alone (**Fig. S3C**). To determine whether this might reflect the fact that the combination induces cytotoxicity, we performed TUNEL staining to evaluate for apoptosis. Indeed, within 12 hours of combined treatment with palbociclib and RMC-5552, we detected a marked increase in apoptotic cells in the combination treatment compared with single agents (**Fig. 3B**, **S3D**).

**Fig 3:**
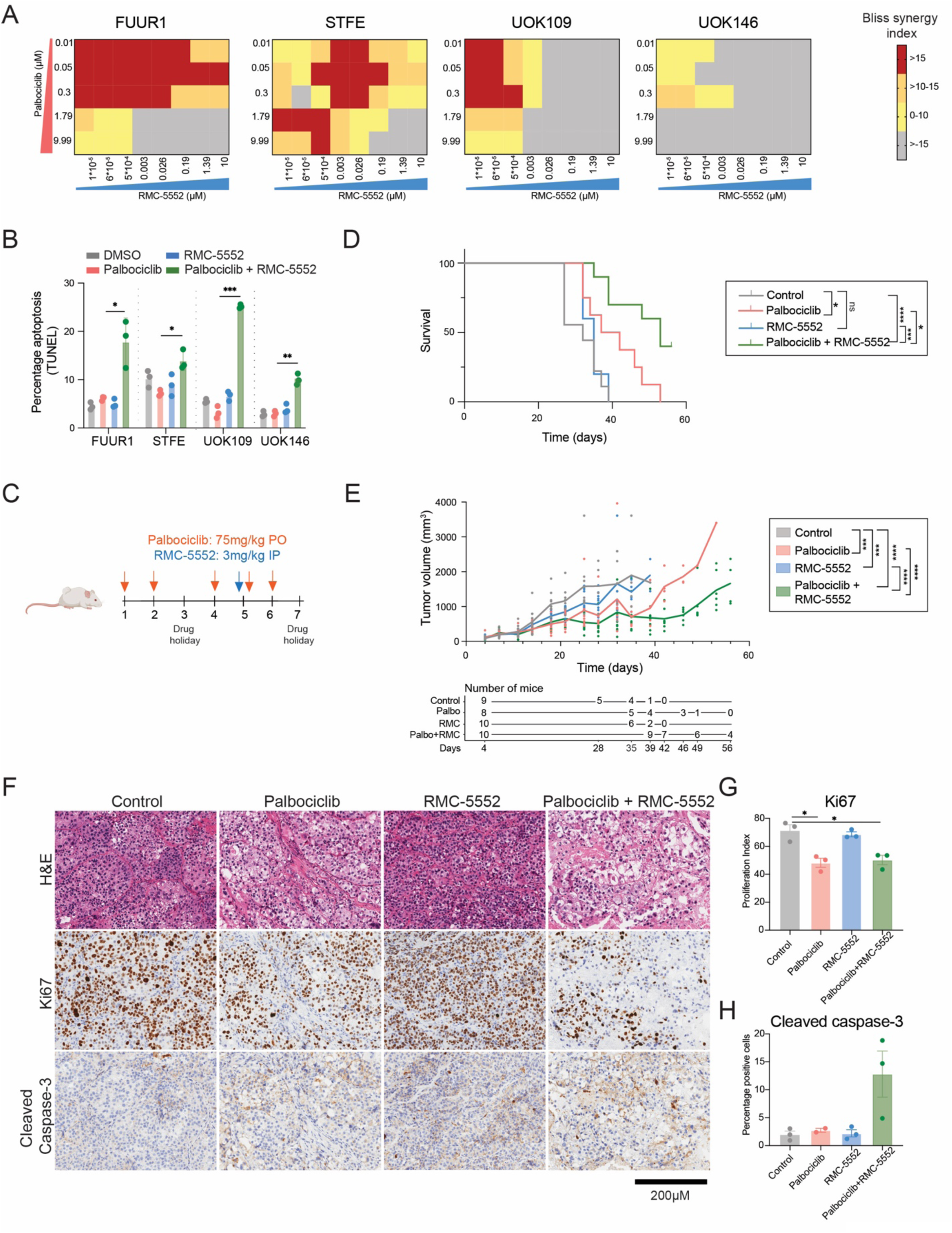
Combined inhibition of CDK4/6 and mTORC1 pathway shows synergy and inhibits tumor growth *in vivo*. (A) Heatmap highlighting Bliss synergy index values for combination doses of palbociclib and RMC-5552 in FUUR1, STFE, UOK109 and UOK146 cell lines, values represent average of three independent biological replicates per cell line. (B) TUNEL assay for apoptosis in cells pre-treated with 300nM palbociclib for 48 hours followed by 15 hours co-treatment with both palbociclib (300nM) and RMC-5552 (200nM) in FUUR1, STFE, UOK109 and UOK146 cell lines, data representative of n=3 biological replicates per cell line, unpaired t-test *p<0.05*, p<0.01**, p<0.001\*\*\** between palbociclib and palbociclib + RMC-5552 groups. (C) Schematic of dosing strategy for FUUR1 xenografts implanted in *NOD.Cg-Prkdc^scid^/J* and treated with a combination of palbociclib and RMC-5552 for a total period of 8 weeks (56 days) or alternative endpoint. (D) Survival curve for individual mice in four arms of the study (starting n per arm: control, n= 9; palbociclib, n=8; RMC-5552, n=10; palbociclib + RMC-5552, n=10). Log-rank test *p<0.05*, p<0.01**, p<0.001\*\*\**. (E) Tumor volume per arm during xenograft study. Individual tumor measurements are shown by dots with mean connected by line. Number of mice in each arm evaluated at measurement time points following survival drops are indicated in the table (*Note*, mice sacrificed in between measurement days are captured in survival curve in D). ANOVA test *p<0.05*, p<0.01**, p<0.001\*\*\**. (F) Histologic images from a representative FUUR1 xenograft tumor in each treatment group (scale bars = 200 μm), upper panel, H&E staining; middle panel, Ki67 staining; lower panel, cleaved caspase-3. (G) Quantification of Ki67 staining across n=3 tumors per arm, unpaired t-test, *p<0.05\**. (H) Quantification of cleaved caspase-3 staining across n=3 tumors for control, RMC-5552, palbociclib + RMC-5552 and n=2 tumors for palbociclib.

Finally, we sought to study the antitumor efficacy of this combination *in vivo.* We performed an initial toxicity study in 8-week old *NOD.Cg-Prkdc^scid^/J* mice to evaluate two doses and frequencies of palbociclib. We observed weight loss with palbociclib at 75 mg/kg given 5 days on/2 days oO with multiple skipped doses required due to weight loss (**Fig. S3E**). However, an altered dosing schedule (3 on/1 oO/2 on/1 oO) of 75 mg/kg palbociclib was well-tolerated in combination with RMC-5552 (3 mg /kg once weekly) (**Fig. S3F**). These doses of both drugs are in line with those previously shown to have activity in solid tumor models with minimal toxicity (51,52).

We then assessed the efficacy of this drug combination in an FUUR1 cell line xenograft model. Animals were treated with placebo, palbociclib (75mg/kg PO 5 times/week), RMC-5552 (3mg/kg IP once/week), or the combination of palbociclib (75mg/kg PO 5 times/week) with RMC-5552(3mg/kg IP once/week). Mice were treated for 8 weeks or until they reached a study-defined endpoint (tumor size or toxicity) (**Fig. 3D**). While RMC-5552 did not have notable single-agent activity *in vivo*, palbociclib led to a period of initial growth suppression, followed by emergence of resistance. However, combination treatment yielded significant tumor growth suppression compared to RMC-5552 or palbociclib alone, with 4 mice surviving at the end of the 8-week study compared with 0 in each of the other three arms (**Fig. 3E**). Importantly, the combination appeared to be well-tolerated with no signs of weight loss or overt toxicity in any of the treatment arms over 8-weeks the treatment period (**Fig. S3G**).

Histological analysis of treated tumors showed that, in comparison to the control tumors, palbociclib and combination treated tumors showed significant reduction in proliferation index as measured by Ki67 index (**Fig 3F-G**), consistent with the cell cycle arrest induced by palbociclib. In addition, combined treatment with RMC-5552 showed a trend toward increase in cleaved caspase-3 activity (*p=0.1* vs. vehicle, unpaired t-test), a marker of apoptotic mediated cell death, in comparison to palbociclib monotherapy, in-line with the observations made *in vitro.* Taken together, the combination of palbociclib with RMC-5552 demonstrated anti-proliferative effects and induced apoptosis *in vivo,* where the combination of palbociclib and RMC-5552 was more effective than palbociclib alone.

## Discussion

Via an integrative genomic approach, we identified CDK4/6 signaling and mTORC1 signaling as highly activated in tRCC. Our findings are consistent with the recent studies characterizing molecular pathways driven by TFE3 fusions in cell line and patient derived xenograft models of tRCC and alveolar soft part sarcoma, a malignancy also driven by a TFE3 fusion (53,54).

In this study, we functionally validated dependence on these pathways through orthogonal genetic and pharmacologic approaches in multiple cell line models of tRCC. We further suggest that the activation of CDK4/6 and mTORC1 pathways in tRCC is intrinsically linked to the pathognomonic driver alteration in tRCC – the TFE3 fusion, a finding that is particularly notable as tRCCs often harbor few other alterations (12,15).

Although inhibition of the CDK4/6 pathway is currently being tested in a variety of solid tumors, it has been most effective in hormone receptor-positive metastatic breast cancer, where CDK4/6 inhibitors have revolutionized the standard of care in combination with ER pathway inhibitors. We suggest that tRCC may represent a promising clinical setting for the broader deployment of CDK4/6i, as the TFE3 fusion is analogous to the ER, with mTORC1 inhibition substituting for ER pathway antagonism. Indeed, our *in vitro* and *in vivo* studies suggest activity to the combination of palbociclib and RMC-5552. Recent work suggests activation of CDK4/6/cyclinD1 signaling in TFE3-fusion driven alveolar soft part sarcoma (ASPS) where inhibition of CDK4/6 activity upon palbociclib treatment induced cell cycle arrest and reduced cell proliferation (54). Our results suggest that the combination of palbociclib with RMC-5552 can further be extended to TFE3-fusion driven ASPS.

While palbociclib has been FDA approved for over a decade, RMC-5552 is currently being tested in Phase I/Ib clinical trials for advanced solid tumors (NCT04774952; NCT05557292). Early data suggest that the drug is well-tolerated with preliminary evidence of activity and plasma exposure at levels consistent with those resulting in p-4EBP1 inhibition in preclinical models (22,55).

Of note, recent preclinical studies have provided a rationale for the combination of HIF2α inhibition and CDK4/6i in clear cell renal cell carcinoma (ccRCC)(56). Moreover, Cyclin D1 has been suggested as a key HIF2α−responsive gene capable of driving resistance to the HIF-2α inhibitor, belzutifan. Therefore, the combination tested in this study may also be applicable to other subtypes of kidney cancer. A recent Phase 1 study of abemaciclib monotherapy in pre-treated metastatic ccRCC showed no single-agent activity (10/11 patients with progressive disease). However, one patient with tRCC (containing clear-cell component histology) was enrolled on this study and was the only patient to experience stable disease (57). The low response rate to abemaciclib in this study may reflect the heavily pre-treated patient population, the requirement for both CDK4 and CDK6 inhibition that would be better achieved with palbociclib, and the typical lack of single-agent activity with CDK4/6i – in addition to the stark molecular differences between ccRCC and tRCC (12,15,35,37).

In conclusion, we demonstrate strong preclinical rationale for inhibition of the CDK4/6 pathway in combination with mTORC1-selective inhibition in tRCC. These data support further clinical evaluation of this therapeutic strategy in tRCC.

## Supplementary Figures

**Figure S1:**
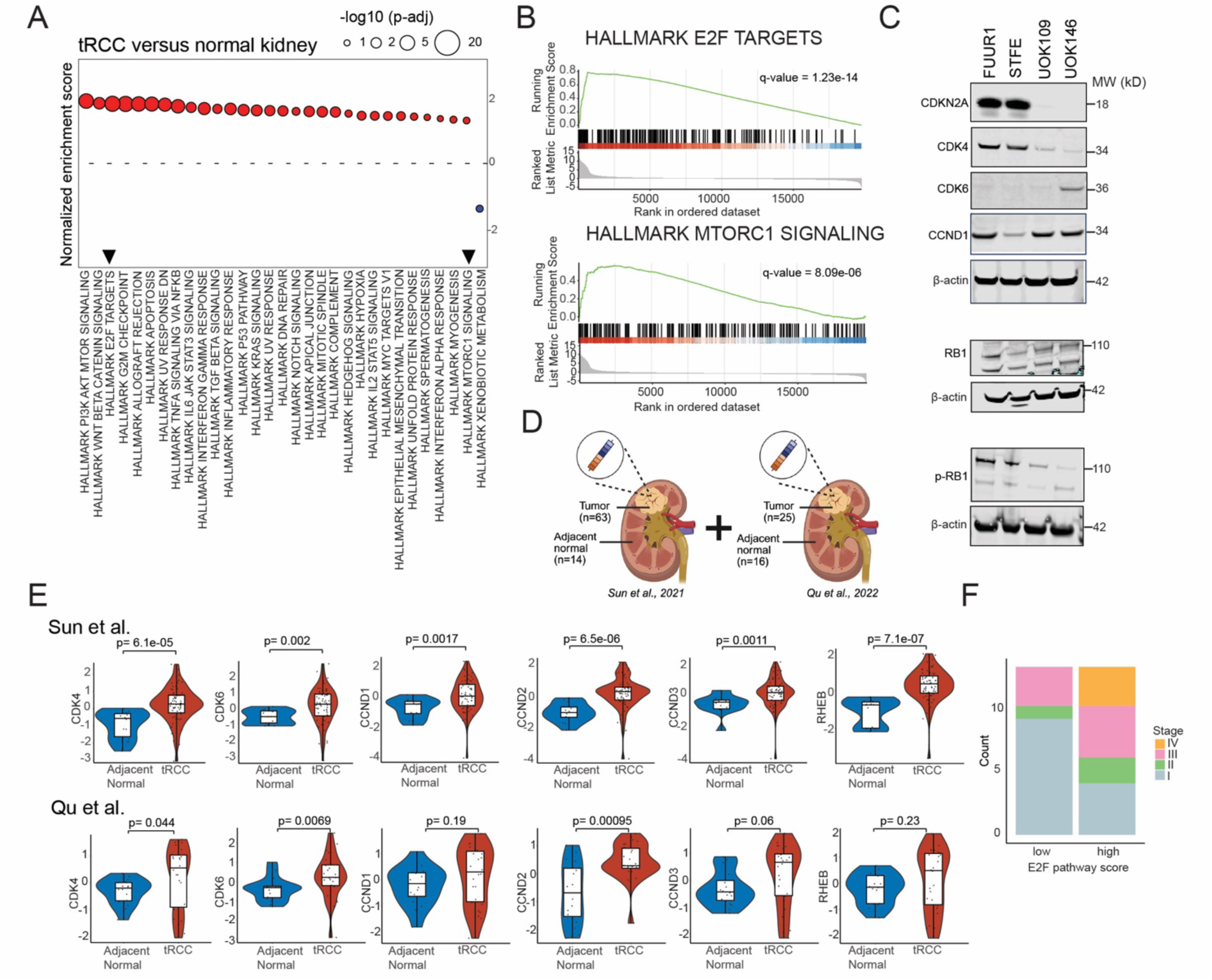
(A) Gene set enrichment analysis for tRCC versus normal subjects in *Qu et al*., 2022 (32), highlighting top 30 enriched pathways (n= 25 tRCC, n= 16 normal). (B) GSEA enrichment plot for E2F targets (top) and mTORC1 signaling (bottom) from analysis in (A) *Qu et al*., 2022 (32). (C) Immunoblot analysis for endogenous protein levels of CDKN2A, CDK4, CDK6, CCND1, phosphor-RB1, RB1 across tRCC cell lines, representative blots for RB1 and phosphor-RB1 were acquired from the same experiment and run on separate gels, data representative of n=3 biological replicates per cell line. (D) Schematic depicting sample collation strategy for both tRCC and normal subjects in *Sun et al*., 2021 (31) and *Qu et al*., 2022 (32). (E) Transcript levels of CDK4, CDK6, CCND1, CCND2 and CCND3, RHEB in tRCC versus normal subjects in *Sun et al*., 2021 (31), unpaired t-test *p<0.05*, p<0.01**, p<0.001\*\*\** (n= 63 tRCC, n= 14 normal) (top), transcript levels of CDK4, CDK6, CCND1, CCND2 and CCND3, RHEB in tRCC versus normal subjects in *Qu et al*., 2022 (32), unpaired t-test *p<0.05*, p<0.01**, p<0.001\*\*\** (n= 25 tRCC, n= 16 normal) (bottom). (F) Stage wise distribution of tRCC patients from *Qu et al., 2022* (*32*), stratified based on *E2F* high and *E2F* low signature based on median value of 1.

**Figure S2:**
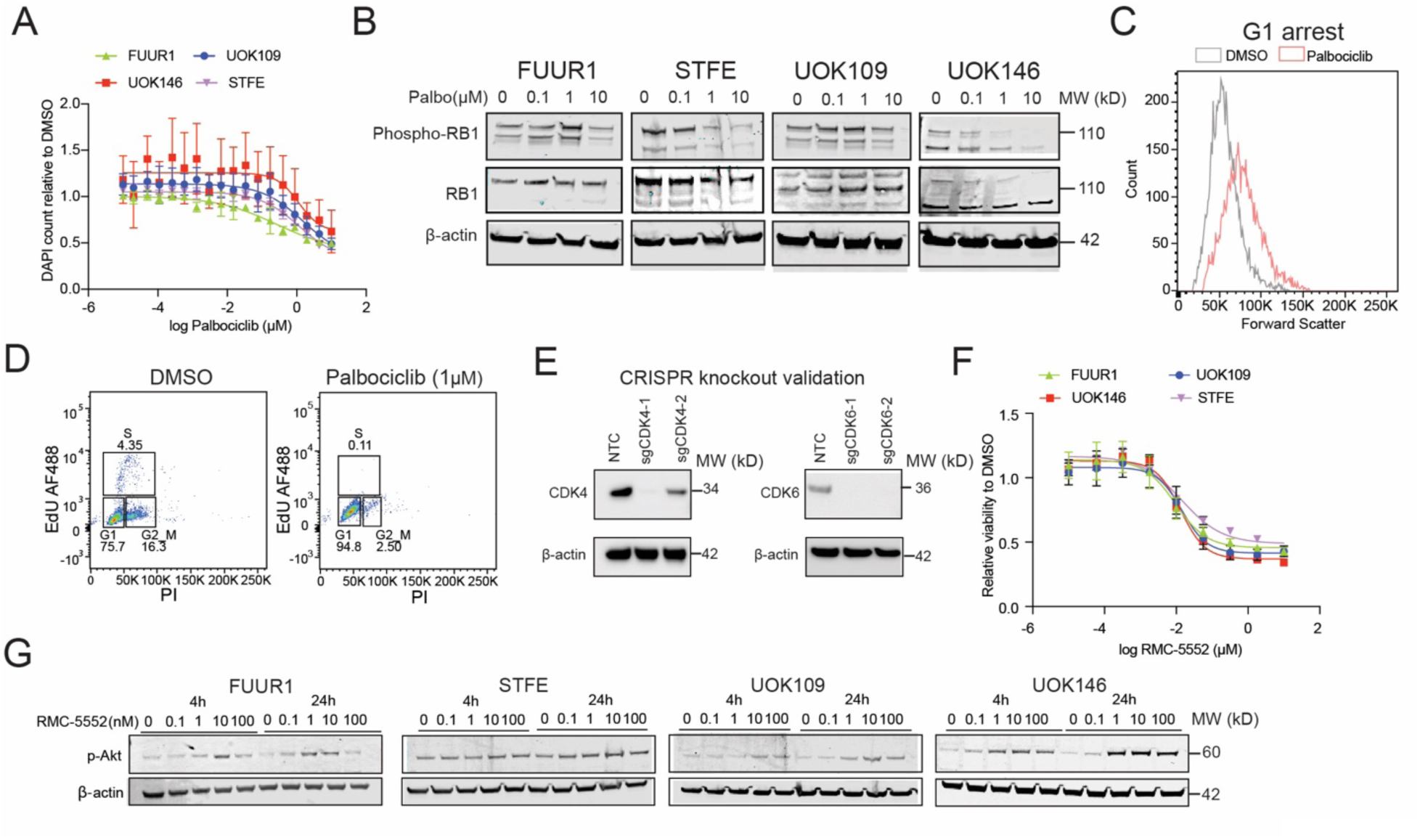
(A) Dose response curve for palbociclib across tRCC cell lines treated within a range of 0.01nM-1mM treated for 72 hours, cell viability was assessed using DAPI nuclei staining with Celigo, IC_50_ values for palbociclib: 250nM-1.5mM, data representative of n=3 biological replicates per cell line. (B) Immunoblot analysis for phospho-Rb and total Rb protein upon palbociclib treatment for 72 hours at different concentrations ranging from 0.01-1µM across 4 tRCC cell lines, data representative of n=3 biological replicates per cell line. (C) Forward scatter profile distribution of UOK146 cells treated with 1mM palbociclib with DMSO as vehicle for 72 hours. (D) Gating strategy for cell cycle distribution of UOK146 cell line treated with 1mM palbociclib (right) with DMSO (left) as vehicle for 72 hours. (E) Immunoblot analysis for endogenous protein levels of CDK4 in FUUR1 and CDK6 in UOK146 upon CRISPR-Cas9 knockout using two different guide RNAs for each gene where NTC is referred as non-targeting control, data representative of n=3 biological replicates per cell line. (F) Dose response curve for RMC-5552 across tRCC cell lines treated within a range of 0.01nM-1mM for 72 hours, cell viability was assessed using Cell Titer Glo assay, IC_50_ value for all cell line are within 10-20nM, for data representative of n=3 biological replicates per cell line. (G) Immunoblot analysis for phospho-Akt, 4 hours and 24 hours post RMC-5552 treatment from 0.1-100nM dose range across tRCC cell lines, data representative of n=3 biological replicates per cell line.

**Figure S3:**
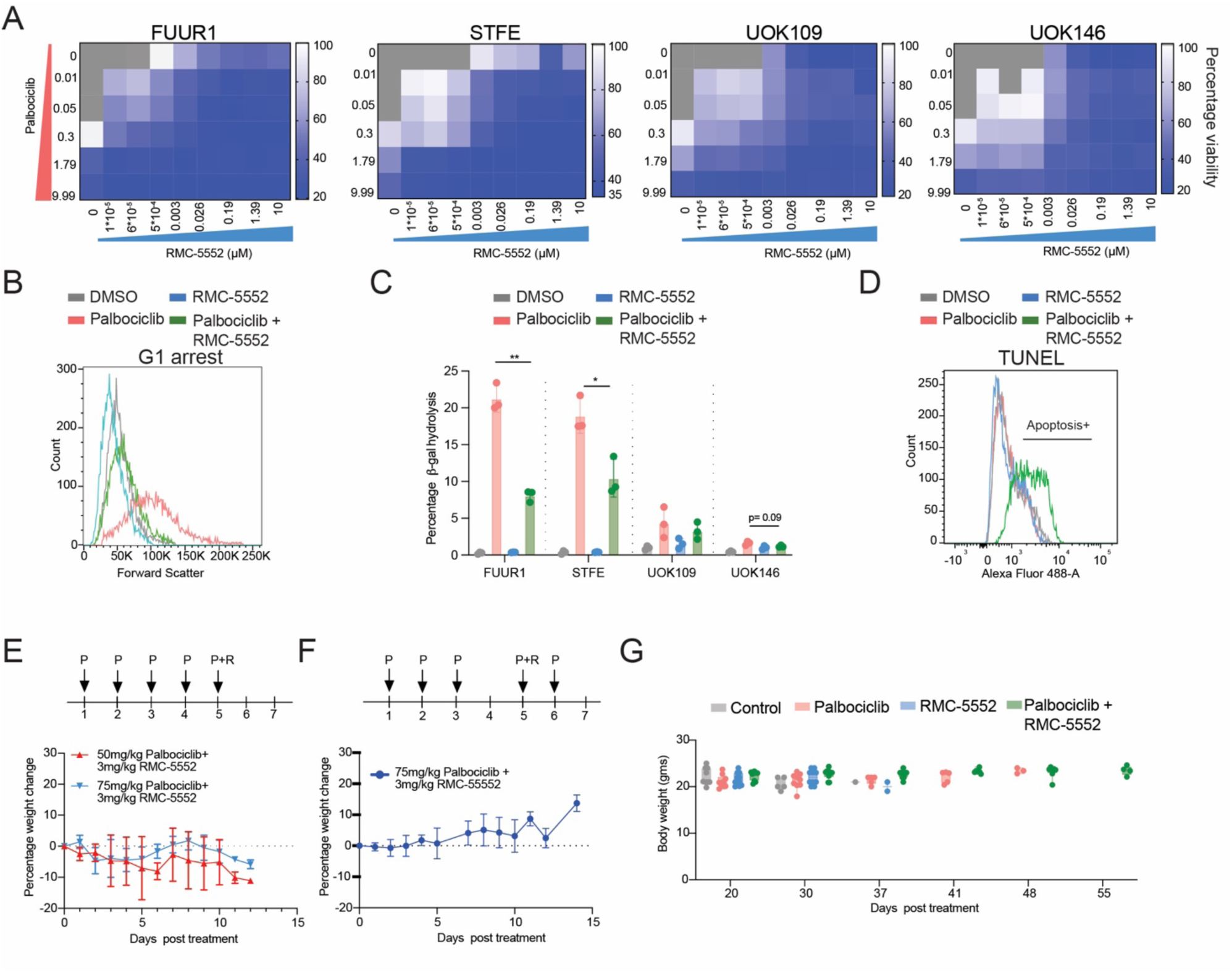
(A) Heatmap of cell viability at combination doses of palbociclib and RMC-5552 calculated using DAPI nuclei staining in FUUR1, STFE, UOK109 and UOK146 cell lines, values represent average of three independent biological replicates per cell line. (B) Forward scatter profile distribution of FUUR1 cells treated with DMSO, 300nM palbociclib, 200nM RMC-5552, and combination of palbociclib with RMC-5552 for 72 hours. (C) Senescence-associated β-galactosidase (SA-β-Gal) activity treated with 300nM palbociclib alone, 200nM RMC-5552 alone, or pre-treated with 300nM palbociclib for 48 hours followed by 72 hours co-treatment with both palbociclib (300nM) and RMC-5552 (200nM), as determined by flow cytometry, data representative of n=3 biological replicates per cell line, unpaired t-test *p<0.05*, p<0.01**, p<0.001\*\*\** between palbociclib and palbociclib + RMC-5552 groups. (D) TUNEL+ labeled cells representative of apoptosis for FUUR1 cell line treated with 300nM palbociclib alone, 200nM RMC-5552 alone, or pre-treated with 300nM palbociclib for 48 hours followed by 15 hours co-treatment with both palbociclib (300nM) and RMC-5552 (200nM), with DMSO as vehicle. (E) Percentage change in weight for *NOD.Cg-Prkdc^scid^/J* mice treated with 50mg/kg palbociclib with 3mg/kg RMC-5552 or with 75mg/kg palbociclib with 3mg/kg RMC-5552 for 5 days consecutively in a week. (F) Percentage change in weight for *NOD.Cg-Prkdc^scid^/J* mice treated with 75mg/kg palbociclib with 3mg/kg RMC-5552 for 5 days / week with one day break as shown. Values represent average of three mice per arm. (G) Weekly mouse body weights for each arm of the experiment from Figure 3.

## Materials and Methods

### RNA-Seq Analysis

RNA-seq data from tRCC and normal adjacent samples were available from two published cohorts (31,32). For the *Sun et al.* cohort, raw gene counts were processed as previously described (31) and available from the supplementary material. TPMs were obtained using the count2tpm function from the “IOBR” R package (v.0.99.8). For the Qu et al. tRCC cohort, RNA-seq FASTQ files were downloaded from NODE (the National Omics Data Encyclopedia, OEP00263). Reads were aligned to the human reference genome (GRCh38.p13 assembly) using STAR (v2.7.10a) and quantified as paired end reads against the GENCODE v45 transcript reference using RSEM (v1.3.1).

Differential gene expression (DGE) was performed within each cohort between tRCC and normal samples using the “DESeq2” package (v.1.40.2) on raw counts. To remove lowly expressed genes, we retained only those genes with fewer than nine zero-count samples and a mean count ≥10 across all samples. Log_2_ fold changes (LFC) were estimated, and p-values were computed using the Wald test. Benjamini-Hochberg correction was used to compute q-values and a q-value<0.05 was taken as statistically significant. Shrunk LFCs, using the apeglm method (“apeglm” R package v.1.22.1) were used for downstream gene ranking.

Gene set enrichment analysis (GSEA) was performed using the Hallmark gene sets (category H) from MSigDB, obtained via the “msigdbr” package (v.7.5.1) for Homo sapiens. The “fgsea” package (v1.26.0) was used to compute normalized enrichment scores (NES) and adjusted p-values for each pathway. Pathways with a Benjamini–Hochberg–adjusted p-value <0.05 were deemed significantly enriched. For pathways of interest, enrichment plots were obtained using the gseaplot2 function from the “enrichplot” R package (v.1.20.0).

The TPM expression of genes of interest (*CCND1, CCND2, CCND3, CDK4, and CDK6*) were log-transformed (log(TPM+1)) and z-score normalized, then compared between tRCC and normal samples using the Wilcoxon rank-sum test. Normalized expression was visualized using the “ComplexHeatmap” R package (v.2.16.0). CDKN2A deletion status was obtained from the corresponding publications as previously described (31,32). For *Qu et al.*, *CDKN2A* loss was defined using a slightly more conservative threshold of −0.3 for the log2 CN ratio (compared to the paper’s 0.1 threshold) to avoid false positives.

### Survival analyses

In the Qu et al. (32) cohort, the activity of the “HALLMARK_E2F_TARGETS” pathway was computed using single-sample GSEA (ssGSEA) (gsva function from the R package “GSVA” (v.1.48.2) with methods = “ssgsea”) on TPM values. E2F pathway activity was dichotomized along the median to define tumors with high vs. low E2F signature. Overall survival (OS) was compared between the 2 groups using the log rank test and visualized using Kaplan-Meier curves (R package “survival” (v.3.5-5) and “survminer” (v.0.4.9.)). Since the tumor staging differed between E2F signature high and low tumors, we performed a multivariable Cox regression model to compare OS between the 2 E2F signature groups while adjusting for stage (defined as early (stages I and II) vs. advanced (III and IV)). R packages, tidyverse” (v.2.0.0), “ggpubr” (v.0.6.0), and “ggsci” (v.3.0.0) were used for data analysis and visualization

### Cell culture

FU-UR-1 (M. Ishiguro’s laboratory, Fukuoka University School of Medicine) and s-TFE (RIKEN, RCB4699), UOK109 (M. Linehan’s laboratory, National Cancer Institute) and UOK146 (M. Linehan’s laboratory, National Cancer Institute), were grown at 37 °C in CO_2_ incubator, in DMEM (Gibco, 11965118) supplemented with 10% FBS (Sigma, F2442), 100 U ml^−1^ penicillin and streptomycin (Gibco, 15140122). Upon confluency, media was removed followed by a rinse with calcium and magnesium free PBS (Gibco, 10010023) and cells were further trypsinized with Trypsin EDTA (Gibco, 25200056).

### Drug Treatment Experiments

All cell lines were seeded in an optically clear bottom 96-well plate (Corning, 3903) at a range of 1000-4000 cells/well. Palbociclib (Selleckchem, PD-0332991) or RMC-5552 (MedChemExpress, HY-132168) both with a stock concentration of 10mM DMSO were added using D300e Digital Dispenser (Tecan) at a dose range of 0.01mM-10mM for Palbociclib and 10pM-10mM for RMC-5552. For single-agent and combination Palbociclib treated cells, 72 hours post treatment, media was removed, cells were rinsed with PBS and fixed with 4% Paraformaldehyde. Nuclei staining was done using Hoescht 33342 (S0485, Selleckchem) and plates were acquired on CeLigo imaging cytometer. For RMC-5552 treated cells, 72 hours post treatment, media was removed, and cell viability was assessed using CellTiter-Glo 2.0 (Promega, G9241). For combination experiments with Palbociclib and RMC-5552, cell viability was assessed using nuclei staining, and the percentage viability was calculated relative to DMSO. Synergy analysis and bliss synergy score were calculated using *Combenefit 2.02* ((58)) at individual combination dose.

### ImmunoBlotting

Cells were resuspended in cell lysis buffer (Cell Signaling Technology, 9803S) supplemented with protease-phosphatase inhibitors (Cell Signaling Technology, 5872S) and given a freeze-thaw cycle to access membrane proteins. Protein quantification was done using Pierce 660nM protein assay (Thermo Fisher Scientific, PI22660) with BSA (Bio-Rad, 5000206) as standard. 40mg of protein for each sample were loaded onto NuPAGE 4–12% Bis-Tris Mini Protein Gels (Thermo Fisher Scientific, NP0335) for separation by SDS–PAGE. Proteins were transferred to PVDF membranes (Life Technologies, IB34002) using an iBlot3 (Thermo Fisher Scientific). Post transfer, blocking was done using Intercept PBS Blocking Buffer (LICOR bio, 927-70001) for 1 hour at room temperature on shaker. Post blocking, membrane was incubated with the indicated primary antibodies at specific dilution in blocking buffer overnight at 4 °C on a shaker. The next day, membranes were washed with PBST three times for 5 minutes each. Post washing, membranes were incubated with secondary antibodies in blocking buffer at 1:5000 (IRDye, LICOR) in blocking buffer on shaker at room temperature. Post secondary incubation, membranes were washed three times with 1X PBST for 5 minutes each and imaged using the Odyssey Classic Imager (LICOR bio).

### Antibodies

CyclinD1 (17H3L3) (ThermoScientific, 701421), p-4EBP1 (Thr37/Thr46) (236B4) (Cell Signaling Technology, 2855), p-4EBP1(Ser65) (174A9) (Cell Signaling Technology, 9456), p-S6 Ribosomal Protein (Ser235/236) (D57.2.2E) (Cell Signaling Technology, 4858), p-Akt (Ser473) (Cell Signaling Technology, 9271), b-actin (C4) (Santa Cruz, sc-47778), Phospho-Rb (Ser807/811) (D20B12) (Cell Signaling Technology, 8566), Rb (4H1) (Cell Signaling Technology, 9309S), CDK4 (DCS-35) (Santa Cruz, SC-23896), CDK6 (DCS83) (Santa Cruz, 3136), CDKN2A p16 INK4A (JC8) (Santa Cruz, 56330)

### CRISPR-Cas9 knockout generation

All sgRNAs were cloned into plentiCRISPRv2 (RRID:Addgene_52961, puromycin resistance) as described (59). All the constructs were confirmed by Sanger sequencing. Lentivirus was prepared by transfecting HEK293T cells with three plasmids: plentiCRISPRv2 (RRID: Addgene_86153), psPAX2 (RRID: Addgene_12260), and pMD2.G (RRID: Addgene_12259) using polyethyleneimine (PEI). Media was replaced with standard growth media after 12 hours, and supernatant containing the virus was collected 48 hours post-transfection. FU-UR-1, s-TFE, UOK109, and UOK146, cell lines were transduced with lentivirus expressing CRISPR-Cas9 and sgRNA targeting the gene of interest, selected by puromycin and blasticidin.

### Cell Cycle Analysis

All cell lines were cultured in a 6-well plate in triplicates and treated with either palbociclib alone from 100nM to 1mM or pre-treated with 300nM palbociclib for 48 hours followed by combination of palbociclib (300nM) and RMC-5552 (200nM) for 72 hours. Post drug treatment, cells were labelled with EdU (20mM) in DMEM for 2 hours at 37 °C in a CO_2_ incubator. Cells were then trypsinized and fluorescently labelled per manufacturer’s instructions (Invitrogen, C10632). DNA content was stained with Propidium Iodide with Ribonuclease A (F10797) and cells were acquired by BD LSR Fortessa. Raw data generated was further analyzed by FlowJo v10.8.

### Cell Growth and Viability Analyses

For cell confluence analysis, all cell lines were seeded in a 6-well plate in triplicates at a density of 0.5-1*10^5^cells per well. 12 hours post seeding, drug containing media was changed with either Palbociclib (300nM) or RMC-5552 (200nM) or DMSO (control) and percentage confluency was measured using CeLigo twice a week. Once the control cells reached maximum confluency, the drug was removed from the medium and treated cells were allowed to grow in the usual culture medium for confluence assessment post drug-washout.

For TUNEL analysis, cell lines were seeded in a 6-well plate in triplicates at a density of 0.5-1*105 cells per well. Cells with combination treatment were pre-treated with palbociclib (300nM) for 48 hours followed by combination of palbociclib (300nM) and RMC-5552 (200nM) for 15 hours. Post treatment, cells were trypsinized, labelled with TdT per manufacturer’s instructions (Roche, 11684795910) and analyzed on BD LSR Fortessa. Raw data generated was further analyzed by FlowJo v10.8.

For senescence analyses, cell lines were cultured in a 6-well plate in triplicates and treated with either palbociclib alone at 1mM or pre-treated with 300nM palbociclib for 48 hours followed by combination of palbociclib (300nM) and RMC-5552 (200nM) for 72 hours. Post drug treatment, cells were trypsinized, fixed and labelled with CellEvent probe per the manufacturer’s conditions (Invitrogen, C10840). Labelled cells were acquired by BD LSR Fortessa and further analyzed by FlowJo v10.8.

For colony forming assays, cell lines were seeded in two 12-well plates at a density of 1000-4000 cells/well and treated with respective drugs or control DMSO, 12 hours post seeding. The cells were maintained under drug treatment conditions for 12 days with media change every third day. After 12 days, cells in one of the plates were fixed with chilled methanol followed by crystal violet staining and cells in other plate were maintained in drug washout conditions for another 12 days. After 24 days, the cells in washout plate were also fixed and stained following same approach. The colonies obtained at both day 12 and 24 were then dissolved in 10% acetic acid and quantified using absorbance at 540nm.

### *In vivo* xenografts

5*10^6^ FU-UR-1 cells were suspended in PBS and Matrigel in a 1:1 ratio and injected subcutaneously into the flanks of 6–8-week-old female mice (SCID mice) from The Jackson Laboratory. In total, 37 mice were injected with 9 mice in control, 8 mice in Palbociclib alone, and 10 mice each in RMC-5552 and combination therapy. Mice were closely observed for tumor formation and the drug treatment was started when tumors became palpable (day 0), with measurements beginning on day 4. Palbociclib was dissolved in an organic solvent with composition 0.5% Tween80, 0.5% methylcellulose and 99% sterile water and was administered five times a week orally at 75mg/kg/dose. RMC-5552 was prepared in 5% Transcutol, 5% Solutol HS-15 and 90% sterile water and was injected intraperitoneally 3mg/kg/dose once a week. Body weights were monitored twice a week, and a dose was skipped if there was more than 10% weight loss. Tumour growth was measured in two perpendicular axes on the indicated days. Tumor volumes (mm^3^) were calculated using the formula (short axis)^2^ × long axis/2. Mice were sacrificed once the tumors reached a threshold of 20 mm in any direction/ showed signs of ulceration or upon weight loss of 15% or more, as per the institutional protocol. The animal study and procedures were performed in accordance with the National Institutes of Health Guide for Care and Use of Laboratory Animals and approved by the Institutional Animal Care and Use Committee of the Dana-Farber Cancer Institute. The mice were housed in groups (five mice per cage) in a 12-h light–dark cycle, with food and water provided *ad libitum*. Ambient temperature (approximately 72 °F) and humidity (approximately 50%) were automatically controlled.

### Statistics and reproducibility

Statistical analyses were performed by GraphPad Prism 10 (v.10.4), Python (on Spyder v.4.1.5), and R (v.4.3.1). No statistical methods were used to predetermine sample sizes. All mice were randomly assigned to different groups. Data collections were not performed blind to the conditions of the experiments. Sample sizes, statistical tests, and significance are described in figure legends. Statistical comparisons were determined by a Mann–Whitney U-test, two-tailed Student’s t-test or ANOVA as indicated in legends. All P values are indicated in the figures and/or legends. All experiments were conducted in replicates as indicated in legends.

## Acknowledgements

We acknowledge Gwo-Shu Mary Lee for helpful discussions regarding the manuscript. Immunohistochemistry was performed at Dana-Farber/Harvard Cancer Center BWH Pathology Cores (P30 CA06516). Schematic illustrations were created with BioRender.com.

## Author contributions

S.R.V designed and supervised the study. T.K.C supervised the study and provided resources. S.G. designed the study and performed the *in vitro* and *in vivo* experiments with assistance from P.Khanna., S.A. and U.A.A. E.S. led the patient data analysis with help from R.M.S., P.Konda., A.S., Q.X. and Z.B. J.L. generated ChIP-seq data analyzed in the study. B.L. and A.S. performed or analyzed CRISPR screening data. W.X. and R.S. contributed to design and interpretation of combination studies. Manuscript, original draft: S.G. and S.R.V. All authors edited and approved the final draft of the manuscript.

## Data availability

External datasets analyzed are public and are available from the respective cited publications.

## Code availability

Algorithms used for data analysis are all publicly available from the indicated references in the paper. No custom code was used in the study.

