## Supplementary figures and images for "Dual Inhibition of CDK4/6 and mTORC1 Establishes a Preclinical Strategy for Translocation Renal Cell Carcinoma"

### Supplementary Figure 1

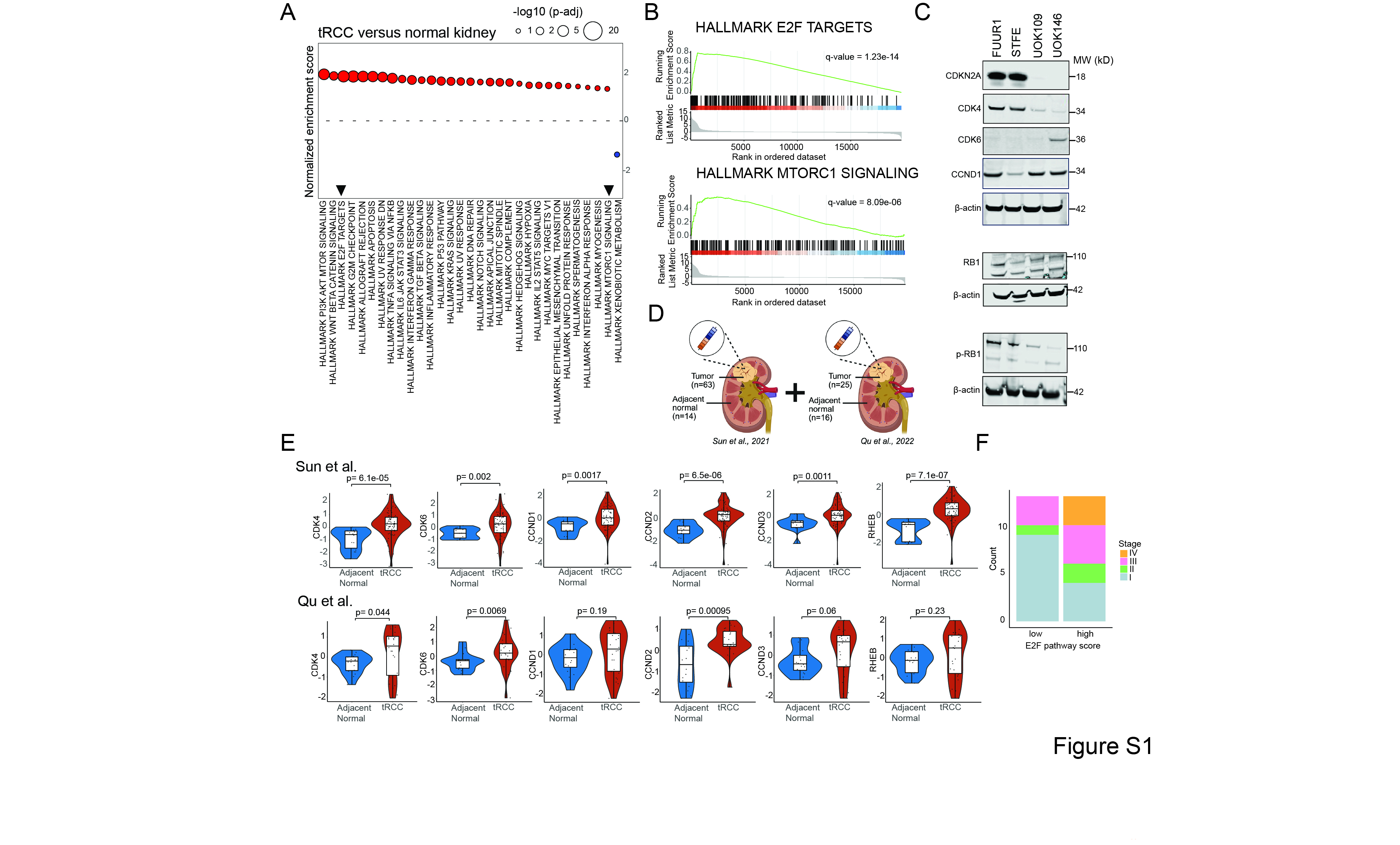

### Supplementary Figure 2

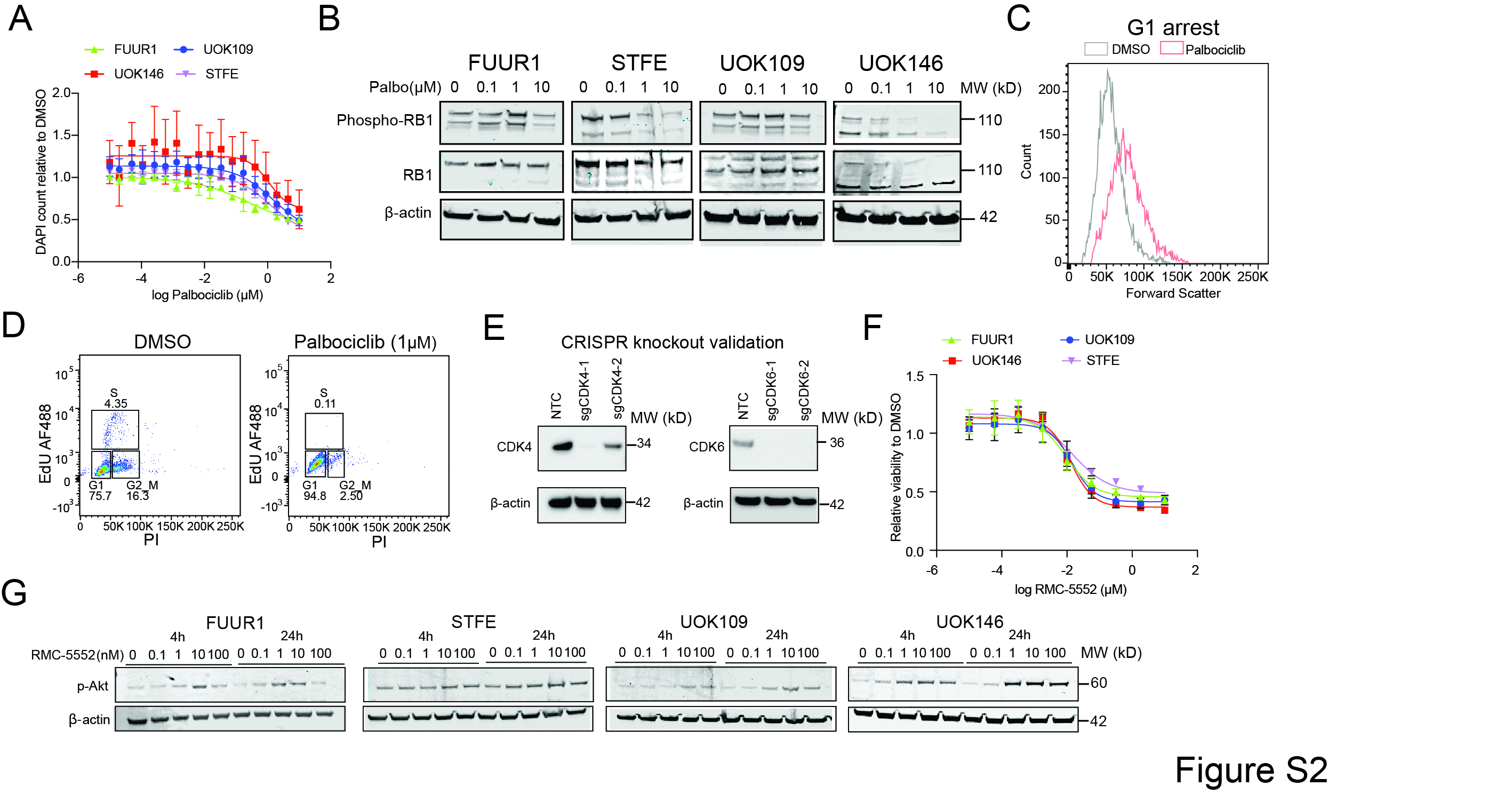

### Supplementary Figure 3

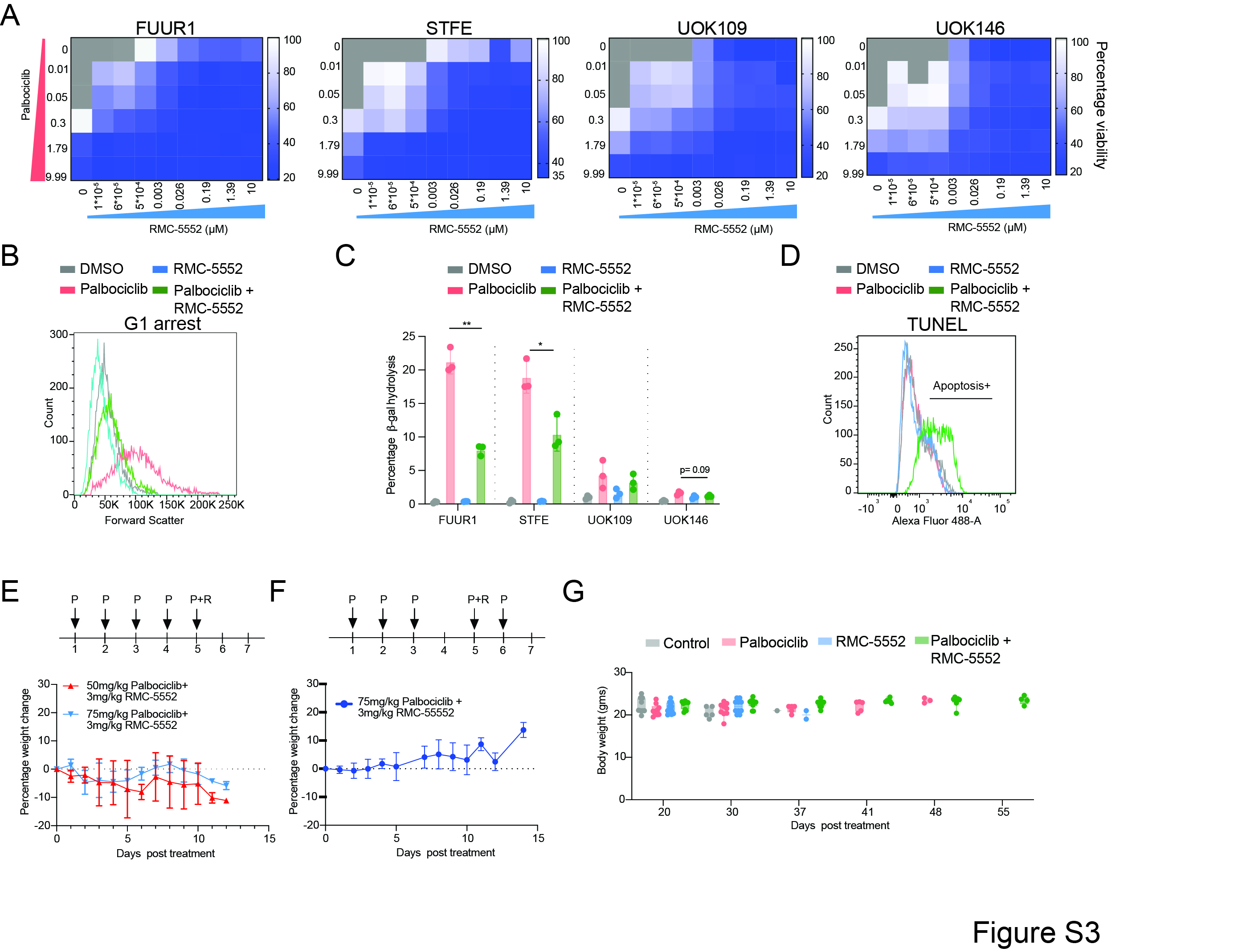
